# De novo Genome Assembly and Annotation of the Seep Tubeworm *Lamellibrachia satsuma* using Oxford Nanopore Sequencing

**DOI:** 10.64898/2026.11.16.767209

**Authors:** Ajit Kumar Patra, Yong Min Kwon, Yong-Jin Won, Youngik Yang

**Author notes:** corresponding author: Youngik Yang.

## Abstract

**Objectives:** The tubeworm *Lamelllibrachia satsuma* inhabits seep region by forming a mutualistic symbiotic relationship with chemosynthetic bacteria residing in its trophosome. Genomic and symbiotic studies have been extensively conducted, primarily from a bacterial perspective. But rarely vice versa. Here we present a high-quality draft genome assembly of *L. satsuma* and it’s tissue-specific transcriptome data. This genome will be valuable for further understanding the mechanisms of adaptation in cold seep environments, host-microbe interactions, as well as genome variation among tubeworms.

**Data description:** We generated ∼103 Gb of genomic sequencing data using Illumina and Oxford Nanopore platforms, which assembled in a 665 Mb draft genome. The assembled genome consisted of 6,320 scaffolds with 95.7% genome completeness. Among 32,394 predicted protein-coding genes, 1,063 gene families have expanded, and 1,385 gene families were contracted. Among the expanded gene families, multiple copies of leukocyte elastase inhibitor, reverse transcriptase, arylesterase, and xanthine dehydrogenase/oxidase genes were found. Molecular clock analysis based on the phylogenomic dataset revealed that *L. sastuma* and *L. luymesi* were diverged from their common ancestor during the Early Cenozoic era, approximately 13 million years ago.

## Objective

Deep-sea hydrothermal vents and cold seeps support diverse marine invertebrates, including siboglinid tubeworms, mussels, and shrimps, that rely on chemosynthetic bacterial symbionts for nutrition[1, 2]. Vestimentiferan tubeworms (Annelida: Siboglinidae) lack a functional digestive system during adulthood and depend entirely on intracellular chemoautotrophic bacteria for their nutritional requirements [3, 4]. These symbionts inhabit a specialized organ, the trophosome, where they synthesize organic compounds through the oxidation of reduced inorganic substances, particularly hydrogen sulfide [5-10].

*L. satsuma*, a vestimentiferan tubeworm originally described from approximately 100 m depth in Kagoshima Bay, Japan [11], inhabits chemosynthetic environments along the Japanese continental margin, including Kagoshima Bay, the Nankai Trough, and the Izu-Ogasawara Ridge [12, 13]. Despite its ecological importance and specialized nutritional symbiosis, genomic resources for *L. satsuma* remain relatively limited. Although a genome assembly and transcriptomic resources have previously been reported by Uchida et al. (2023)[14], independently generated genomic and tissue-specific transcriptomic datasets can provide complementary resources for comparative genomic analyses, functional gene characterization, and further investigations into host–microbe interactions and molecular adaptations to chemosynthetic environments.

The primary objective of this study was to generate and characterize a draft reference genome and tissue-specific transcriptomic datasets for *L. satsuma*. To achieve this, genomic DNA extracted from vestimentum tissue was sequenced using Illumina NovaSeq 6000 and Oxford Nanopore GridION platforms. Additionally, RNA sequencing was performed on vestimentum and trophosome tissues to provide transcriptional evidence for genome annotation and facilitate future investigations of tissue-specific gene expression.

The resulting genomic and transcriptomic datasets, including a 665-Mb draft genome and its structural and functional annotations, are presented as publicly accessible resources. These data are intended to support future comparative genomic, evolutionary, and functional studies of vestimentiferan tubeworms, particularly those investigating genomic adaptations to chemosynthetic environments and the molecular basis of host–symbiont associations.

## Data description

### Sample collection and sequencing

*L. satsuma* specimens were collected from Kagoshima Bay, Japan, at approximately 110 m depth, as previously described [15]. Following removal of the chitinous tubes, tissues were washed with 70% ethanol, rinsed with sterile seawater, and stored at −80°C. Genomic DNA was extracted from vestimentum tissue using the phenol–chloroform method.

A 550-bp paired-end genomic library was prepared using the TruSeq DNA PCR-Free kit and 2⍰× ⍰151⍰bp reads were sequenced on an Illumina NovaSeq 6000, generating 53.25 Gb of raw data (Table 1). Oxford Nanopore libraries were prepared using the SQK-LSK109 ligation kit and sequenced on a GridION X5, producing 50.03 Gb of long reads. Basecalling was performed using Guppy in high-accuracy mode (Table 1).

**Table 1:** Overview of data files/data sets.

| **Label** | **Name of data file/data set** | **File type** | **Data repository and identifier** |
|---|---|---|---|
| Data set 1 | Illumina NovaSeq 6000 whole-genome paired-end sequencing reads | FASTQ (.fastq.gz) | NCBI SRA: SRR15602513 |
| Data set 2 | Oxford Nanopore GridION whole-genome long-read sequencing data | FASTQ (.fastq.gz) | NCBI SRA: SRR15602504–SRR15602512 |
| Data set 3 | Illumina HiSeq 2000 vestimentum RNA-seq paired-end reads | FASTQ (.fastq.gz) | NCBI SRA: SRR2641679 |
| Data set 4 | Illumina HiSeq 2000 trophosome RNA-seq paired-end reads | FASTQ (.fastq.gz) | NCBI SRA: SRR2641680 |
| Data set 5 | Genome assembly | FASTA (.fna) | NCBI Accession: JAHXPS000000000 |
| Data set 6 | Assembly summary | Table file (.xlsx) | Figshare: **https://doi.org/10.6084/m9.figshare.35016802** |
| Data set 7 | BUSCO scores | Table file (.xlsx) |  |
| Data set 8 | Statistics of repetitive elements | Table file (.xlsx) |  |
| Data set 9 | Genome annotation | Table file (.xlsx) |  |
| Data set 10 | Rapidly expanded gene families | Table file (.xlsx) |  |
| Data set 11 | Time-calibrated phylogenetic tree showing evolutionary divergence | Figure file (.png) |  |

Vestimentum and trophosome tissues were preserved in RNAlater and stored at -20°C. For RNA extraction, tissues were homogenized under liquid nitrogen and processed using TRIzol. RNA quality was assessed using an Agilent 2100 Bioanalyzer. Following messenger RNA enrichment and cDNA library preparation, Illumina HiSeq 2000 sequencing generated 4.95 Gb and 4.79 Gb of transcriptomic data from vestimentum and trophosome, respectively (Table 1).

### Sequence processing and genome assembly

Illumina reads were quality-filtered using Trimmomatic [16], retaining 47.60 Gb. Nanopore adapters were removed using PoreChop, retaining 49.82 Gb. Nanopore reads exceeding 1 kb (43.45 Gb) were assembled using Flye [17]. Draft assemblies were polished using Racon [18] and Medaka, followed by two rounds of Pilon [19] correction with Illumina reads. Following removal of nine contigs flagged during database submission, the final genome assembly comprised 664.98 Mb distributed across 6,320 contigs, with an N50 of 267.06 kb (Supplementary Table 1) and 95.7% completeness assessed using BUSCO metazoa_odb10 database (Supplementary Table 2).

### Repeat identification and gene annotation

Repetitive elements were identified using Tandem Repeats Finder [20], RepeatModeler [21], and RepeatMasker [22]. Repetitive sequences constituted 62.89% of the genome, including tandem repeats (17.61%), unclassified repeats (16.77%), and long interspersed nuclear elements (14.07%). Identified repeats were masked before gene prediction (Supplementary Table 3).

Protein-coding genes were predicted using BRAKER [23], incorporating GeneMark-ET [24], AUGUSTUS [25], and RNA-seq alignments generated using TopHat [26]. Homology-based predictions employed TBLASTN [27], GenBlastA [28], and Exonerate [29], with additional protein evidence from Swiss-Prot [30]. Conflicting and unsupported gene models were excluded.

The final annotation comprised 32,394 protein-coding genes. Functional annotations were assigned using DIAMOND [31] against UniRef90 and InterProScan [32] for protein domains and Gene Ontology terms. Additionally, 761 transfer RNA genes and 29 ribosomal RNA genes were identified using tRNAscan-SE and RNAmmer, respectively (Supplementary Table 4).

### Comparative genomic analysis

Orthologous gene families were identified using OrthoMCL [33]. A dataset of 194 single-copy orthologs was aligned using MUSCLE [34] and trimmed using trimAl [35]. Phylogenetic reconstruction was performed using RAxML [36] with the PROTGAMMAJTT model and 100 bootstrap replicates. Divergence times were estimated using MEGA [37], applying a 753-million-year calibration between Caenorhabditis elegans and Octopus vulgaris obtained from TimeTree [38] (Supplementary Fig.1). Gene family analyses using CAFE [39] identified 1,063 expanded and 1,385 contracted gene families (Supplementary Table 5).

## Limitations

The reference is a draft, contig-level assembly rather than a chromosome-level assembly. The available tissue transcriptomes represent two tissue types and the supplied manuscript does not document biological replicate numbers. Gene predictions and functional annotations are computational and require independent validation for specific biological applications.

## Abbreviations

BUSCO: Benchmarking Universal Single-Copy Orthologs
NCBI: National Center for Biotechnology Information
RNA-seq: RNA sequencing
SRA: Sequence Read Archive

## Declarations

### Availability of data and materials

The genomic and transcriptomic sequencing datasets generated in this study have been deposited in the NCBI Sequence Read Archive (SRA). Oxford Nanopore long-read sequencing data are available under accession numbers SRR15602504–SRR15602512, Illumina paired-end whole-genome sequencing data under SRR15602513, and Illumina RNA-seq data from vestimentum and trophosome tissues under SRR2641679 and SRR2641680, respectively. The draft genome assembly of *L. satsuma* is publicly available in NCBI GenBank under accession number JAHXPS000000000. Additional datasets, including genome assembly statistics, BUSCO completeness assessments, repetitive element statistics, genome annotation, rapidly expanded gene families, and a time-calibrated phylogenetic tree, have been deposited in Figshare and are publicly accessible at https://doi.org/10.6084/m9.figshare.35016802.

### Authors’ contributions

Y.Y. conceived concept and designed the experiments, A.K.P. performed experiments, Y.Y. conducted analyses, and Y.Y. wrote the paper. All authors reviewed the manuscript.

## Acknowledgements

We would like to thank the captain and crew of R/V Natsushima and the operation team of ROV Hyper-Dolphin, Japan Agency for Marine-Earth Science & Technology (JAMSTEC) for collecting and providing samples for this study.

---

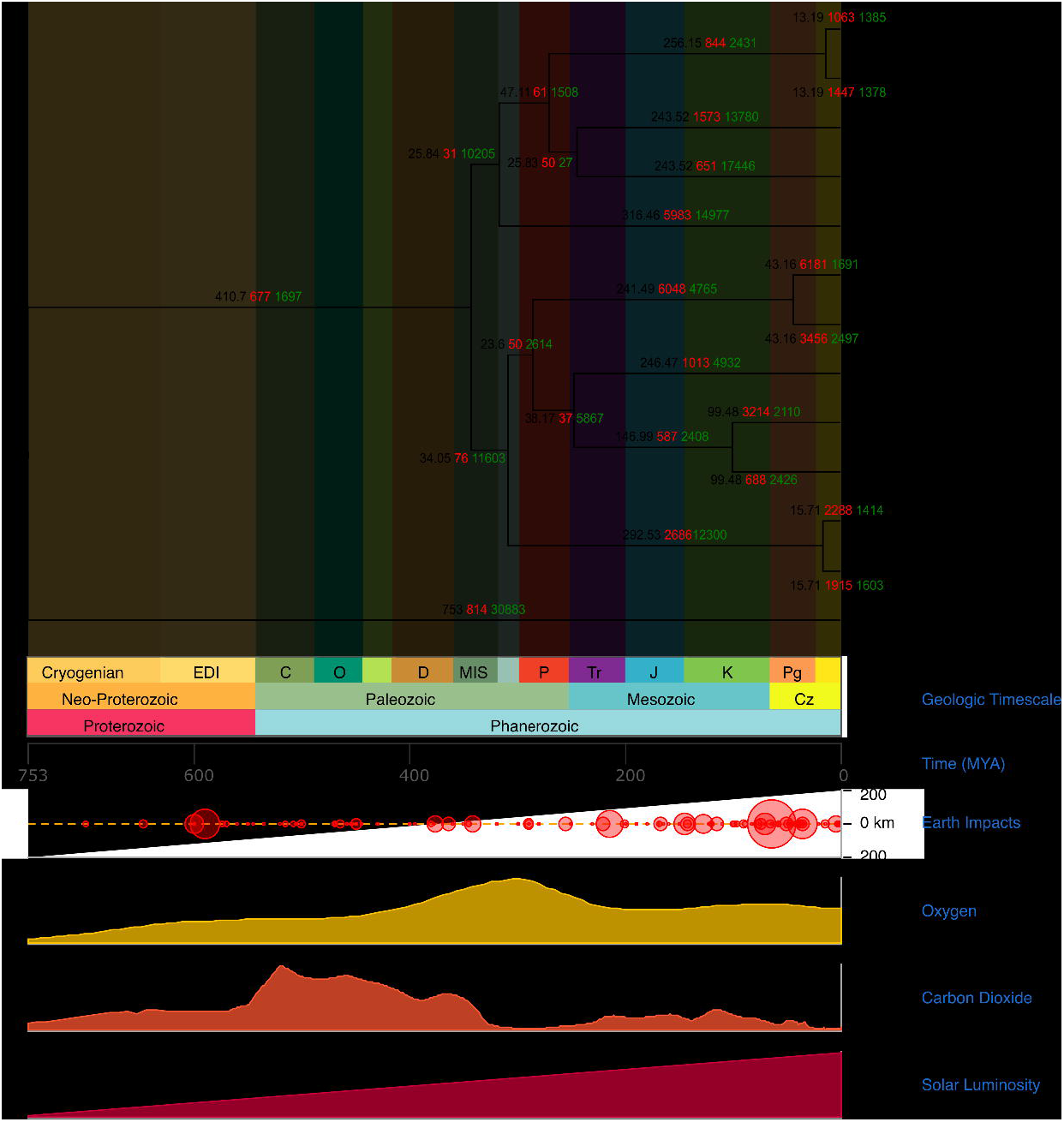

